# Simulation of a sudden drop-off in distal dense core vesicle concentration in *Drosophila* type II motoneuron terminals

**DOI:** 10.1101/2021.03.04.434010

**Authors:** I. A. Kuznetsov, A. V. Kuznetsov

**Affiliations:** Perelman School of Medicine, University of Pennsylvania, Philadelphia, PA 19104, USA; Department of Bioengineering, University of Pennsylvania, Philadelphia, PA 19104, USA; Department of Mechanical and Aerospace Engineering, North Carolina State University, Raleigh, NC 27695-7910, USA

**Keywords:** neurons, axonal transport, neuropeptide transport, dense core vesicles, mathematical modeling

## Abstract

Recent experimental observations have shown evidence of an unexpected sudden drop-off in the dense core vesicles (DCVs) content at the ends of certain types of axon endings. This paper seeks to determine whether these observations may be explained without modifying the parameters characterizing the ability of distal *en passant* boutons to capture and accumulate DCVs. We developed a mathematical model that is based on the conservation of captured and transiting DCVs in boutons. The model consists of 77 ordinary differential equations and is solved using a standard Matlab solver. We hypothesize that the drop in DCV content in distal boutons is due to an insufficient supply of anterogradely moving DCVs coming from the soma. As anterogradely moving DCVs are captured (and eventually destroyed) in more proximal boutons on their way to the end of the terminal, the fluxes of anterogradely moving DCVs between the boutons become increasingly smaller, and the most distal boutons are left without DCVs. We tested this hypothesis by modifying the flux of DCVs entering the terminal and found that the number of most distal boutons left unfilled increases if the DCV flux entering the terminal is decreased. The number of anterogradely moving DCVs in the axon can be increased either by the release of a portion of captured DCVs into the anterograde component or by an increase of the anterograde DCV flux into the terminal. This increase could lead to having enough anterogradely moving DCVs such that they could reach the most distal bouton and then turn around by changing molecular motors that propel them. The model suggests that this could result in an increased concentration of resident DCVs in distal boutons beginning with bouton 2 (the most distal is bouton 1). This is because in distal boutons, DCVs have a larger chance to be captured from the transiting state as they pass the boutons moving anterogradely and then again as they pass the same boutons moving retrogradely.

## 1. Introduction

Recent research suggests that many neurodegenerative diseases, such as Charcot-Marie-Tooth disease [1], hereditary spastic paraplegia [2], spinal muscular atrophy [2], Parkinson’s disease (PD) [3], and Alzheimer’s disease [4-6] are associated with defects of axonal transport. Various mathematical approaches have been developed to analyze the links between neurodegeneration and axonal transport defects. Examples of such research include modeling of the pileups of axonal cargos [7], formation of amyloid plaques and its effect on axonal transport [8], development of traffic jams after traumatic brain injury [9], prion-like features of neurodegenerative diseases [10], as well the effects of tau protein on regulating anterograde and retrograde transport in axons [11].

Neuropeptides are one of many substances actively transported in the axon. They are important for regulating mood, motivation, sleep, and drug addiction [12,13]. Neuropeptides’ synthesis occurs in the soma. Following synthesis, neuropeptides are transported in dense core vesicles (DCVs) along the length of the axon toward the axon terminals (Fig. 1a) [14]. Kinesin-1 and kinesin-3 motors drive anterograde transport of DCVs while cytoplasmic dynein motors drive retrograde transport [15]. Dynein can also contribute to the anterograde transport of vesicles that are associated with small microtubules (MTs) by moving small MTs along non-motile large MTs [16,17].

**Fig. 1.**
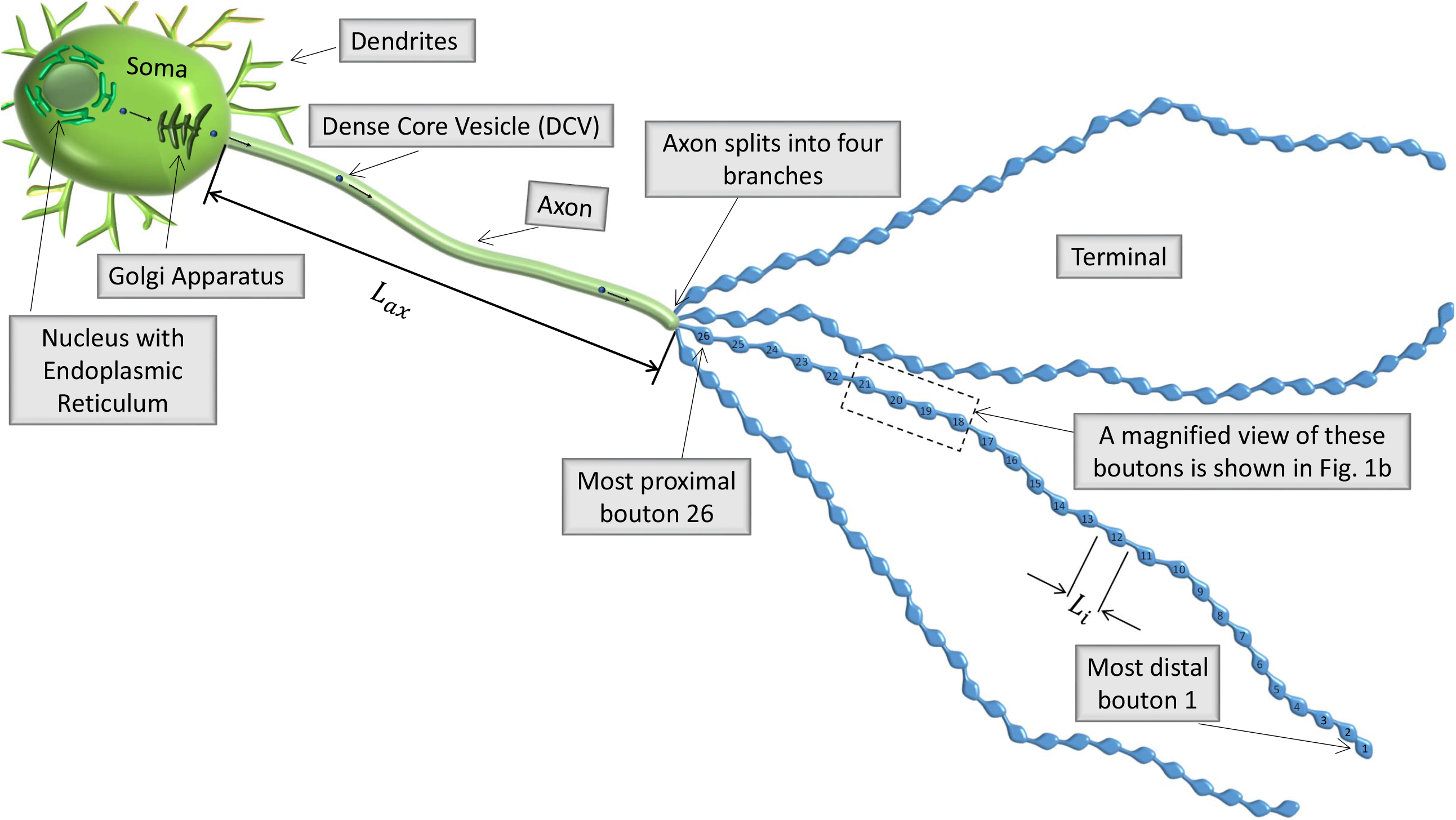
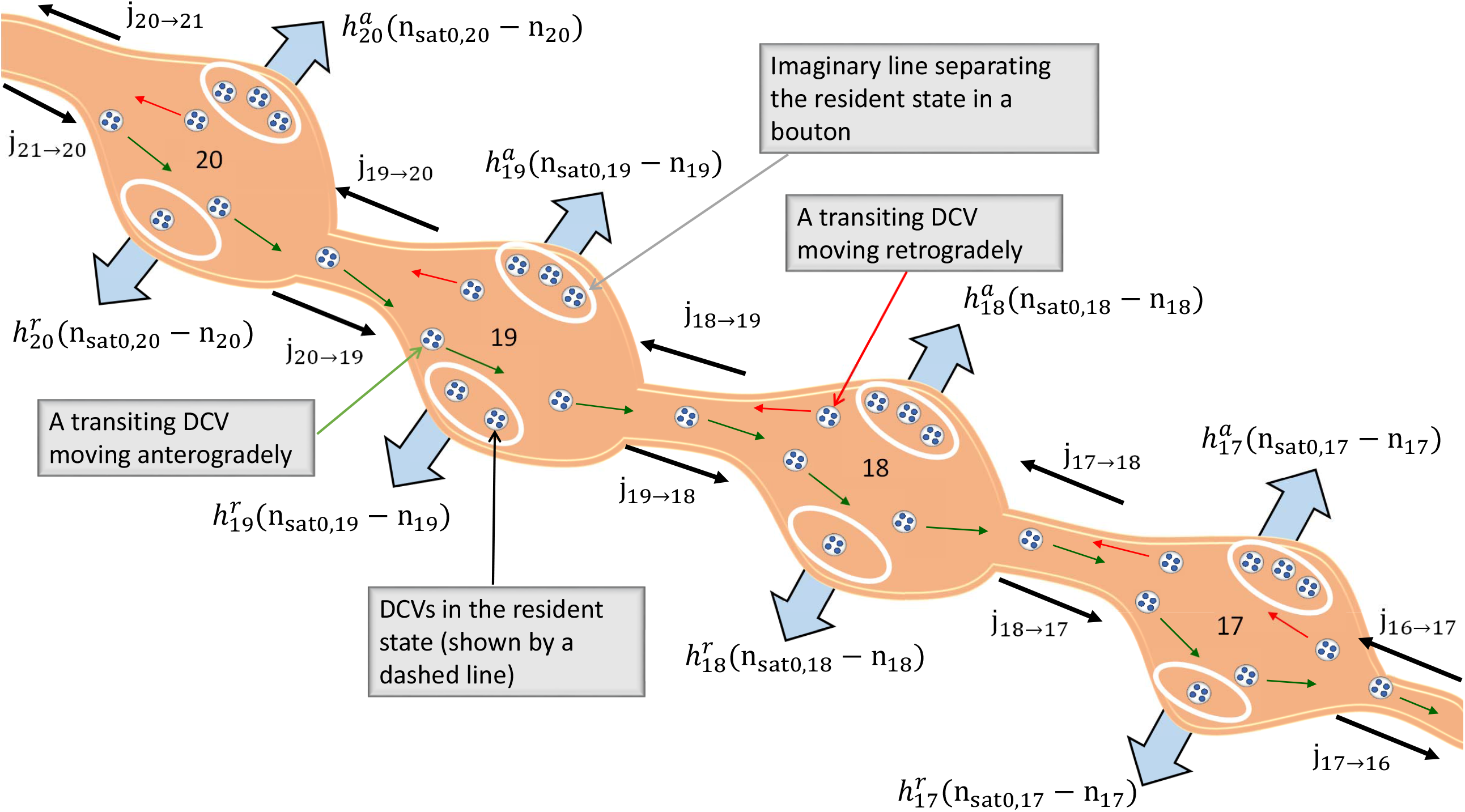
(a) A schematic diagram of a neuron with an axon; the arbor of the axon splits into four identical branches. We assumed that each branch contains 26 boutons. Boutons are numbered consistent with the convention used in [20], starting with the most distal (#1) to the most proximal (#26). (b) A magnified portion of the terminal showing boutons 20, 19, 18, and 17. Transiting and resident DCVs are depicted, as well as DCV fluxes between the boutons. Block arrows illustrate the capture of transiting DCVs into the resident state. The rates of DCV capture are also shown.

DCV transport in axons of *Drosophila* motoneurons with type I, II, and III endings was investigated in [18-21]. These axonal endings are characterized by different morphologies; they have different numbers and sizes of *en passant* boutons (hereafter boutons). Boutons are sites that accumulate and release neurotransmitters; they appear as varicosities located along the length of the axon terminals. Models of DCV transport and capture in axon terminals were developed in [22-27].

In *Drosophila*, type II terminals have the largest number of boutons. In some branches of type II terminals ref. [18] recorded a drop-off in the DCV content that unexpectedly appeared at the farthest ends of the arbor. This striking finding merits an investigation on why this unintuitive feature emerges. It should be noted that many branches do not show the drop-off, and if one thinks about the innervation of different muscles by a single axon as a tree structure, the drop-off usually appears in the most distal branches. Ref. [27] demonstrated that the drop-off could be explained by the assumption that approximately 20% of the most distal boutons are characterized by different parameter values. The novelty of the present paper is the demonstration that the appearance of the drop-off zone can be explained without the assumption that the most distal boutons have any special properties, specifically, that they have more limited capacity for DCVs than other boutons. Our motivation is to explain the drop-off by the dynamics of transport processes rather than by assigning a special set of parameters to most distal boutons.

In this paper, we investigate whether problems with the supply of distal boutons with various organelles, which are typical, for example, in PD, may result from instability in organelle transport. One can envision that PD can be associated with instability in axonal transport, which leads to an insufficient supply of various organelles at the presynaptic terminals and other distal active sites. Although our research does not directly simulate processes leading to the development of PD in a human brain, the neurons that produce type II boutons in *Drosophila* and the neurons that die in humans with Parkinson’s disease (PD) have two things in common: they are monoaminergic, which means they are sensitive to autoxidation, and they have large axonal arbors [18,28]. Thus a study of neurons with type II terminals may be useful to better understand PD.

Here we demonstrate that the depletion of active sites (boutons) in the distal axon can be caused by an imbalance of anterograde and retrograde transport. This approach may lead to furthering our fundamental understanding of underlying reasons for neurodegenerative diseases.

Following experimental observations [29], we assumed that DCV content is constant until the sudden drop-off zone starts. We do not pre-suppose the drop-off zone in the model, but the model can predict this zone if simulations show that the flux of anterogradely transported DCVs is depleted before DCVs reach distal boutons.

## 2. Materials and models

### 2.1. Governing equations

We followed the model developed in [27] and investigated whether this model can predict a drop-off zone of DCVs at the end of the terminal without assuming any special properties for the most distal boutons.

A terminal with 26 boutons was simulated (Fig. 1a). The value of 26 comes from a previous estimate, presented in [18], of the percentage of anterogradely moving DCVs that reach the most distal bouton. The model simulates DCV concentrations in resident, anterograde transiting, and retrograde transiting kinetic states in boutons (Fig. 2). Following the convention adopted in [20], the boutons are numbered from the most distal #1 to the most proximal #26 (Fig. 1a). DCV concentrations in resident and transiting states were characterized by DCV linear number density which describes the number of DCVs per unit length in the terminal. We used a multi-compartment model [30-32] to formulate the conservation equations, with the number of DCVs in a compartment being a conserved quantity.

**Fig. 2.**
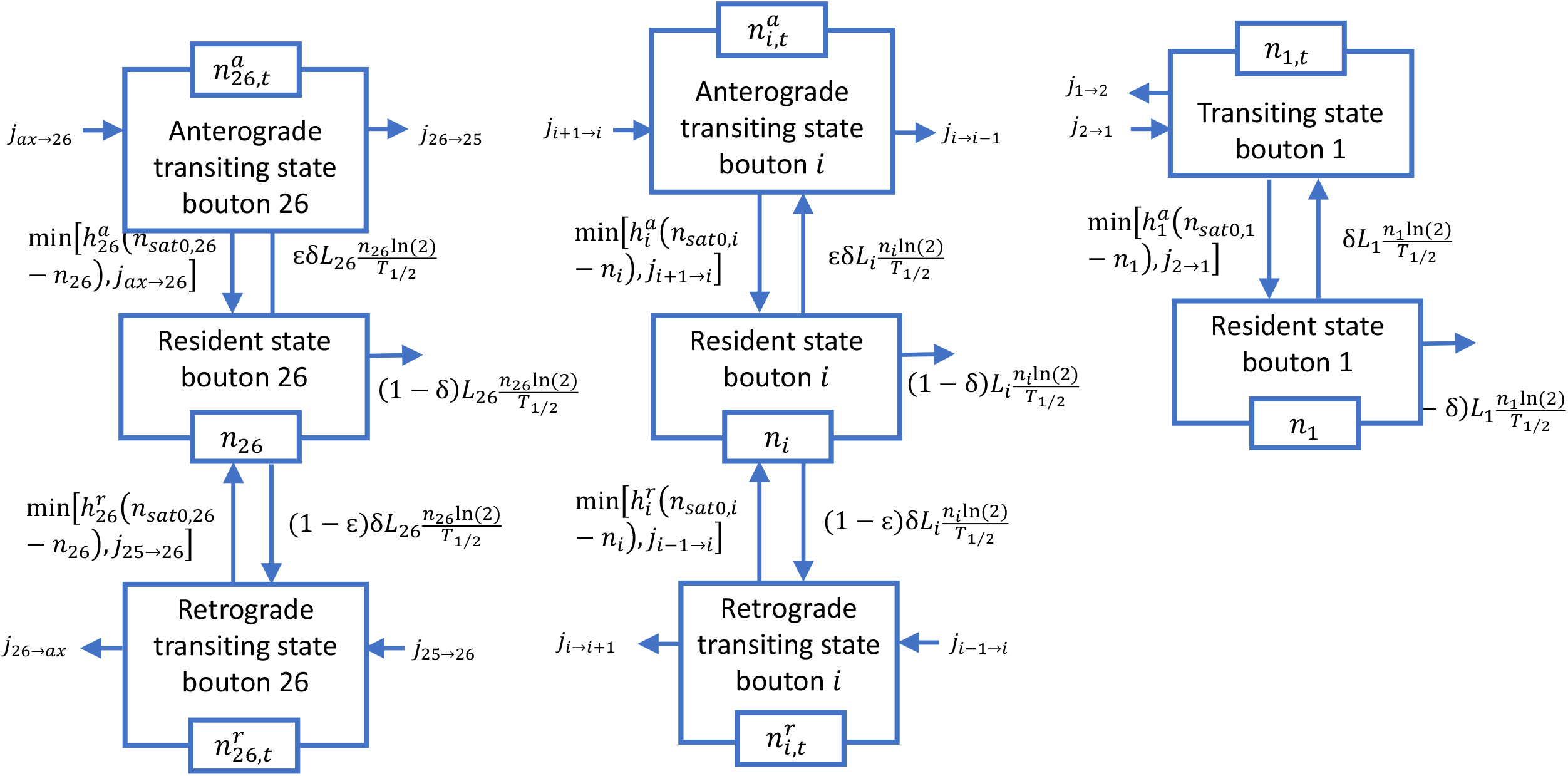
A diagram of a compartmental model illustrating transport in the transiting and resident states in the terminal. Arrows display DCV exchange between the transiting states in adjacent boutons, DCV capture into the resident state, and re-release from the resident state. DCV destruction in the resident state is also shown.

Except for bouton 1, all boutons are represented by three compartments: the compartment containing resident DCVs, the compartment containing transiting DCVs that move anterogradely, and the compartment containing transiting DCVs that move retrogradely (Fig. 2). Separate compartments are introduced for anterograde and retrograde transiting DCVs to avoid mixing of DCVs re-released from the resident state with DCVs already residing in the transiting state. Without this separation, it would be impossible to assign a certain portion of re-released DCVs (*ε*) to the anterograde component and the remainder (1 − *ε*) to the retrograde component. Bouton 1 is represented by only two compartments: the one containing resident DCVs and the other containing transiting DCVs. This is because DCVs turn around in bouton 1, changing the direction of their motion from anterograde to retrograde.

The following equations express the conservation of DCVs in the most proximal bouton (Fig. 2):

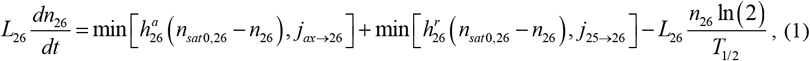

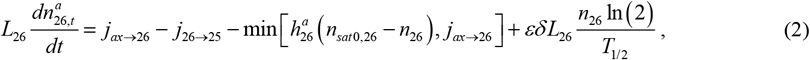

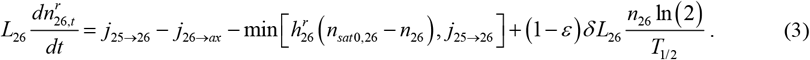

Eq. (1) describes the conservation of resident DCVs in bouton 26, Eq. (2) describes the conservation of transiting DCVs moving anterogradely in bouton 26, and Eq. (3) describes the conservation of transiting DCVs moving retrogradely in bouton 26. The terms on the left-hand sides of Eqs. (1)-(3) represent the change in the number of DCVs in the corresponding kinetic state, the terms involving *j* on the right-hand sides of Eqs. (2) and (3) represent the net DCV transfer into the transiting states in boutons (DCV inflow minus DCV outflow), the terms involving “min” on the right-hand sides of Eqs. (1)-(3) represent the DCV capture by the resident state and the corresponding DCV loss by the transiting states, and the terms involving *T*_1/2_ in Eqs. (1)-(3) represent re-release of captured DCVs from the resident state into the transiting states. In Eqs. (1)-(3) *n*, 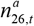, and 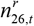 are the concentrations of resident, anterograde transiting, and retrograde transiting DCVs, respectively, in bouton *i* (*i*=1,…, 26); *t* is the time; *n*_*sat*0,*I*_ is the saturated concentration of resident DCVs in bouton *i* at infinite DCV half-life or at infinite DCV residence time (*i*=1,…, 26); *L*_*i*_ is the length of a compartment occupied by bouton *i* (defined in Fig. 1a, *i*=1,…, 26); 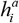 and 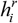 are the mass transfer coefficients characterizing the rates of DCVs capture into the resident state in bouton *i* (2,…, 26) as DCVs pass through bouton *i* in the anterograde and retrograde directions, respectively; *j*_*ax*→26_ is the anterograde flux of new DCVs from the axon to the most proximal bouton (#26); *j*_*j* →*k*_ is the flux of DCVs from compartment “*j*” to compartment “*k*” (see Fig. 1b); *j*_26→*ax*_ is the retrograde flux of DCVs from the most proximal bouton back to the axon; and *T*_1/2_ is the half-life or half-residence time of DCVs captured into the resident state in boutons. In terms of DCV conservation in the resident states in boutons, it does not matter whether DCVs are destroyed or released back to the transiting pool. If DCVs are destroyed, *T*_1/2_ is interpreted as the half-life. If DCVs are released back to circulation, *T*_1/2_ is interpreted as the half-residence time. The parameter *δ* defines what happens to DCVs that are captured into the bouton’s resident state: *δ* = 0 implies that all DCVs are eventually destroyed in boutons (or that the DCV content leaves the boutons by spontaneous exocytotic events) while *δ* = 1 implies that DCVs, after spending some time in the resident state, are released back to the transiting pool. Also, the parameter *ε* indicates how DCVs released from the resident state are split between the anterograde and retrograde components: *ε* is the portion of released DCVs that join the anterograde component and (1 − *ε*) is the portion of DCVs that join the retrograde component. The term min 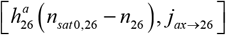 on the right-hand side of Eq. (1) simulates the fact that the rate at which a bouton captures anterogradely moving DCVs cannot exceed the flux of anterogradely moving DCVs into the bouton. A similar role is played by the term min 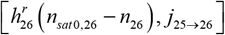

We note that in our model we use the term release as a term that opposes capture. In biology, release also refers to releasing the contents inside of vesicles to the extracellular milieu by exocytosis. Contrary to this, in our model we consider DCV release as returning DCVs back into the transiting DCV pool.

Equations that define the conservation of DCVs in boutons 25 through 2 are (Fig. 2)

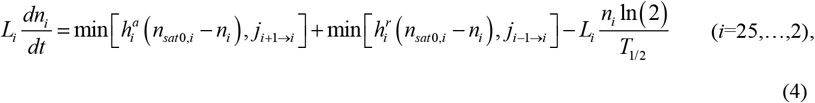

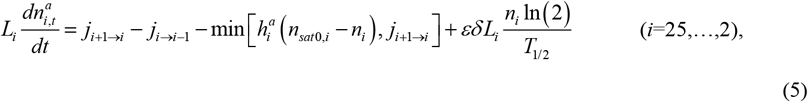

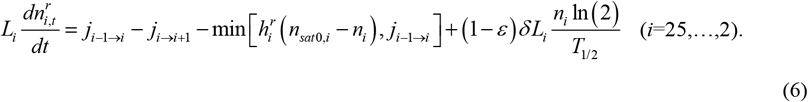

Eq. (4) describes the conservation of resident DCVs in boutons 25 through 2, Eq. (5) describes the conservation of anterograde transiting DCVs in boutons 25 through 2, and Eq. (6) describes the conservation of retrograde transiting DCVs in these boutons.

Equations that describe the conservation of DCVs in the most distal bouton are (Fig. 2)

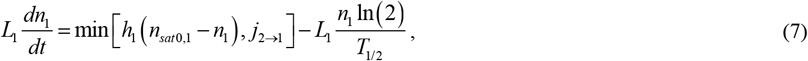

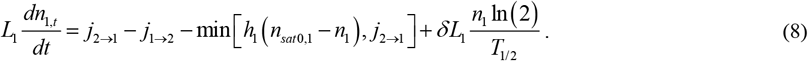

Eq. (7) describes the conservation of resident DCVs in bouton 1 and Eq. (8) describes the conservation of transiting DCVs in this bouton. DCVs pass bouton 1 only once, and thus there are no separate anterograde and retrograde transiting states in this bouton. In Eqs. (1) and (2), *h*_1_ is the mass transfer coefficient characterizing the rate of DCV capture into the resident state in bouton 1 (DCVs pass bouton 1 only once) and *n*_1,*t*_ is the concentration of transiting DCVs in bouton 1.

Eqs. (1)-(8) include DCV fluxes between various boutons and between the most proximal bouton and the terminal. These fluxes need to be modeled. In this paper, we assumed that *j*_*ax*→ 26_ remains constant during the process of filling the terminal. The DCV fluxes have units of vesicles/s. The anterograde flux of transiting DCVs between the most proximal bouton (bouton 26) and bouton 25 is simulated by the following equation:

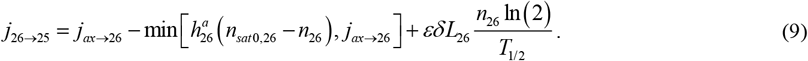

The following equations were used for modeling anterograde fluxes between boutons 25 through 2:

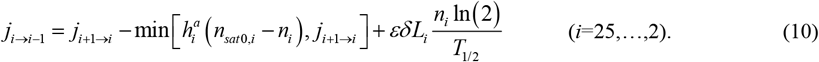

The retrograde flux from bouton 1 into bouton 2 was modeled by the following equation:

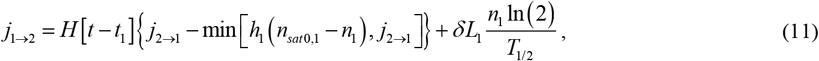

where *H* is the Heaviside step function and *t*_1_ is the time required for DCVs to change the direction in the most distal bouton if they are not captured. The Heaviside step function delays the onset of the retrograde flux of DCVs by the time *t*_1_, avoiding a physically contradictory situation when the retrograde DCV flux would start simultaneously with the anterograde flux. Ideally, the delay should be applied to all DCVs arriving into the most distal bouton (not only to the initial ones), but this does not seem to be possible in the framework of the compartmental model. The initial delay is expected to have a minor effect on filling the terminal because *t*_1_ = 300 s (Table 1) is small relative to how long it takes to fill the boutons with DCVs (hours), see Table 4.

**Table 1.** Model parameters that were estimated based on values found in the literature or assumed on physical grounds.

| Symbol | Definition | Units | Estimated value(s) or range | Reference(s) |
| --- | --- | --- | --- | --- |
| $b$ | Parameter characterizing an increase of DCV capture efficiency from one bouton to the next more distal bouton (boutons #$i$ to #$i-1$) | | 1.08<sup>a</sup> | |
| $L_i$ | Length of a compartment occupied by bouton $i$ ($i = 1, \dots, 26$), as depicted in Fig. 1a | $\mu\text{m}$ | 10 | [29] |
| $n_{0,t}$ | Concentration of DCVs in the transiting state at $t = 0$ | vesicles/<br>$\mu\text{m}$ | 1 | |
| $t_1$ | Average time it takes for DCVs in the most distal bouton to change the direction of motion, if they are not captured (probably involves the change of molecular motors that propel DCVs) | s | 300 | [20,29] |
| $T_{1/2}$ | Half-life or half-residence time of DCVs in the resident state in boutons | s | $2.16 \times 10^4$ | [19] |
| $\varepsilon$ | The portion of DCVs released from the resident state that join the anterograde component; $(1 - \varepsilon)$ is the portion of released DCVs that join the retrograde component. | | 0-1 | |
| $\delta$ | The portion of DCVs released from the captured state back to the transiting state; $(1 - \delta)$ is the portion of captured DCVs that are eventually destroyed in boutons. | | 0-1 | |
| $\varpi$ | DCV capture efficiency in the most proximal bouton, defined as the percentage of DCVs captured in an empty bouton when DCVs pass the bouton. | | 0.1 | [18] |

<sup>a</sup> This value is selected as the minimum value for which $h_1$ remains positive.

Retrograde fluxes between boutons 2 through 25 were modeled by utilizing the following equations:

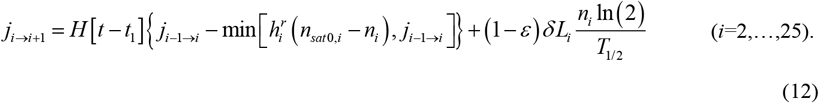

We modeled the retrograde flux that leaves the terminal from the most proximal bouton by the following equation:

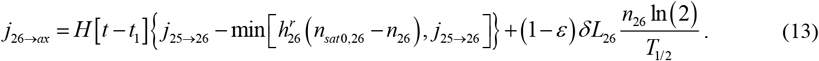

Eqs. (1)-(8) define a system of 77 ordinary differential equations (ODEs) of the first order; 77 initial conditions are therefore needed.

We assumed that initially the resident state in the terminal was empty:

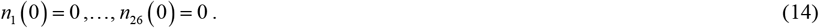

We also assumed that the initial DCV concentration was constant and uniform in the transiting states:

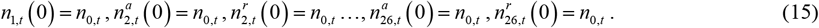

### 2.2. Estimation of parameter values that are involved in the model

#### 2.2.1. Two groups of parameters involved in the model, divided by the method that was used to estimate them

We divided the parameters of the model into two groups. We estimated the parameters belonging to the first group (*b, L*_1_, …, *L*_26_, *L*_*ax*_, *n*_0,*t*_, *t*_1_, *T*_1/2_, *ε, δ*, and *ϖ*) utilizing data found in published literature or assumed on physical grounds. We summarize these parameters in Table 1. We estimated the values of the parameters belonging to the second group (*n*_*sat,ax*_, *n*_*sat*, 1_, …, *n*_*sat*, 26_, *h*_1_, …, *h*_26_, *h*_*in*_, *n*_*sat*0,1_, …, *n*_*sat*0,26_, *n*_*sat,ax*_, and *j*_*ax*→26_) by stating the conservation of DCVs at *t* = 0 and at steady-state. We summarized these parameters in Tables 2 and 3. Below, we explain how the parameter values belonging to the second group were calculated.

**Table 2.** Mass transfer coefficients that characterize the rates of DCV capture into the resident state; the units of all *h*_*i*_ in Table 2 are μm/s.

| $h_1$ | $h_2$ | $h_3$ | $h_4$ | $h_5$ | $h_6$ | $h_7$ |
| --- | --- | --- | --- | --- | --- | --- |
| $2.62 \times 10^{-5}$ | $3.62 \times 10^{-5}$ | $4.65 \times 10^{-5}$ | $5.71 \times 10^{-5}$ | $6.80 \times 10^{-5}$ | $7.92 \times 10^{-5}$ | $9.07 \times 10^{-5}$ |

| $h_8$ | $h_9$ | $h_{10}$ | $h_{11}$ | $h_{12}$ | $h_{13}$ | $h_{14}$ |
| --- | --- | --- | --- | --- | --- | --- |
| $1.03 \times 10^{-4}$ | $1.15 \times 10^{-4}$ | $1.27 \times 10^{-4}$ | $1.40 \times 10^{-4}$ | $1.53 \times 10^{-4}$ | $1.67 \times 10^{-4}$ | $1.81 \times 10^{-4}$ |

| $h_{15}$ | $h_{16}$ | $h_{17}$ | $h_{18}$ | $h_{19}$ | $h_{20}$ | $h_{21}$ |
| --- | --- | --- | --- | --- | --- | --- |
| $1.96 \times 10^{-4}$ | $2.11 \times 10^{-4}$ | $2.26 \times 10^{-4}$ | $2.42 \times 10^{-4}$ | $2.58 \times 10^{-4}$ | $2.74 \times 10^{-4}$ | $2.92 \times 10^{-4}$ |

| $h_{22}$ | $h_{23}$ | $h_{24}$ | $h_{25}$ | $h_{26}$ |
| --- | --- | --- | --- | --- |
| $3.09 \times 10^{-4}$ | $3.27 \times 10^{-4}$ | $3.46 \times 10^{-4}$ | $3.65 \times 10^{-4}$ | $3.85 \times 10^{-4}$ |

**Table 3.**
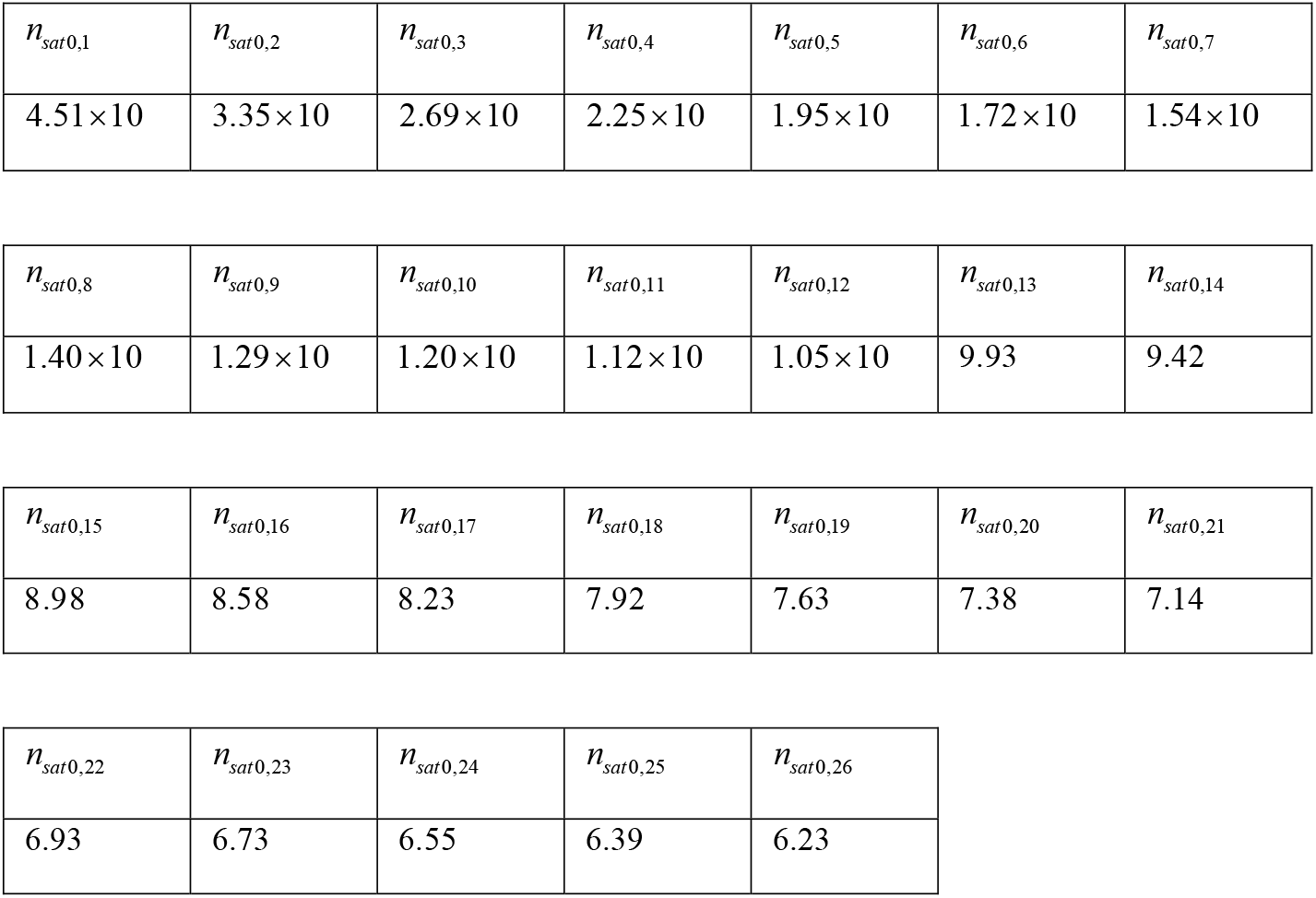
Saturated concentrations of DCVs in boutons at infinite DCV half-life or at infinite DCV residence time; the units of *n*_*sat*0,*I*_ in Table 3 are vesicles/μm.

|  |  |  |  |  |  |  |
| --- | --- | --- | --- | --- | --- | --- |
| $n_{sat0,1}$ | $n_{sat0,2}$ | $n_{sat0,3}$ | $n_{sat0,4}$ | $n_{sat0,5}$ | $n_{sat0,6}$ | $n_{sat0,7}$ |
| $4.51 \times 10$ | $3.35 \times 10$ | $2.69 \times 10$ | $2.25 \times 10$ | $1.95 \times 10$ | $1.72 \times 10$ | $1.54 \times 10$ |
| $n_{sat0,8}$ | $n_{sat0,9}$ | $n_{sat0,10}$ | $n_{sat0,11}$ | $n_{sat0,12}$ | $n_{sat0,13}$ | $n_{sat0,14}$ |
| $1.40 \times 10$ | $1.29 \times 10$ | $1.20 \times 10$ | $1.12 \times 10$ | $1.05 \times 10$ | 9.93 | 9.42 |
| $n_{sat0,15}$ | $n_{sat0,16}$ | $n_{sat0,17}$ | $n_{sat0,18}$ | $n_{sat0,19}$ | $n_{sat0,20}$ | $n_{sat0,21}$ |
| 8.98 | 8.58 | 8.23 | 7.92 | 7.63 | 7.38 | 7.14 |

|  |  |  |  |  |
| --- | --- | --- | --- | --- |
| $n_{sat0,22}$ | $n_{sat0,23}$ | $n_{sat0,24}$ | $n_{sat0,25}$ | $n_{sat0,26}$ |
| 6.93 | 6.73 | 6.55 | 6.39 | 6.23 |

#### 2.2.2. Estimation of saturated concentrations of DCVs in the resident state in boutons, *n*_*sat, I*_ (*i* = 1,…, 26)

Type II terminals, which were investigated in [18], were considered. These terminals have about 80 boutons per muscle, spread over 3 or 4 branches [29]. Based on this data, we estimated that there are approximately 26 boutons per single branch. When a type II bouton is filled to saturation, it contains approximately 34 DCVs [18]. We assumed the following uniform steady-state distribution of DCVs in different boutons to model this situation:

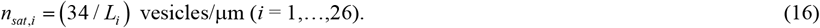

It should be noted that Eq. (16) gives an estimate of the saturated DCV concentration, and it is used only in estimating model parameters in section 2.2.3. Ref. [18] reported that the drop-off in the DCV content appears suddenly at the farthest ends of the arbor. Therefore, Eq. (16) may not correctly approximate the steady-state distribution in the most distal boutons, if the drop-off occurs. However, we do not want to postulate any distinction between the boutons in the model, since we are interested in simulating the drop-off based on the physics of DCV transport.

#### 2.2.3. Estimation of mass transfer coefficients characterizing capture of DCVs into the resident state in boutons, *h*_1_, *h*_2_, …, *h*_26_ ; saturated concentrations of DCVs in boutons at infinite DCV half-life or half-residence time, *n*_*sat*0,1_, *n*_*sat*0,2_, …, *n*_*sat*0,26_; and a DCV flux from the axon into the most proximal bouton, *j*_*ax*→26_

Mass transfer coefficients that characterize DCV capture as the DCVs pass the boutons in anterograde and retrograde directions were assumed to be equal:

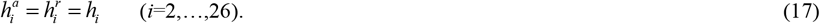

Two types of statements were used to estimate the values of *h*_*i*_ and *n*_*sat*0,*I*_ (*i*=1,…, 26). The DCV capture rate at *t* = 0 in various boutons was first stated:

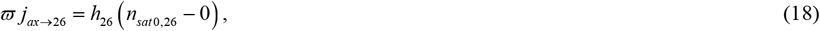

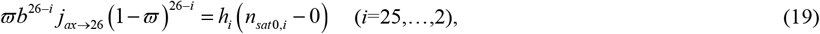

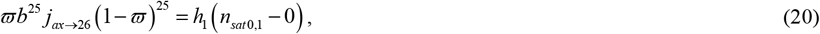

where *ϖ* is the capture efficiency, which is a parameter characterizing the percentage of captured DCVs as they pass an empty bouton [18,26]. More distal boutons are hungrier (*b* = 1.08); our model shows that the values of *h*_*i*_ decrease from the most proximal to the most distal bouton (Table 2), and 1.08 is the minimum value for which *h*_1_ remains positive.

The second type of statements deals with the rate of DCV capture at steady-state. If the resident DCVs’ half-life or half-residence time is finite, the capture must still proceed, even at a steady state. Here we are most interested in investigating the situation when at steady-state all DCVs are captured while moving anterogradely before they reached the most distal bouton:

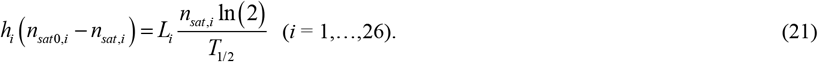

The value of the DCV flux entering the most distal bouton from the axon is not assumed, but is computed by the following equation:

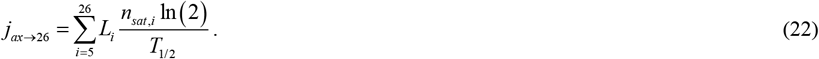

We used Matlab’s (Matlab R2019a, MathWorks, Natick, MA, USA) solver Solve; the results are summarized in Tables 2 and 3.

From Eq. (22) it follows that *j*_*ax*→26_= 2.40 ×10^−2^ vesicles/s, which is in the range of 3.33×10^−2^ vesicles/s reported in [18,20].

## 3. Results

The system of 77 ODEs described in section 2.1 was solved by Matlab’s solver, ODE45 (Matlab R2019a, MathWorks, Natick, MA, USA). The model predicts a drop in the concentration of resident DCVs in the four most distal boutons (Fig. 3a). The reason for this is the reduction in anterograde DCV fluxes between the boutons (Fig. 3b). Thus the numerical results show the drop-off of the DCV concentration reported in [18]. The drop-off was not postulated *a priori* in our model (see Eq. (16)), it is predicted by the model.

**Fig. 3.**
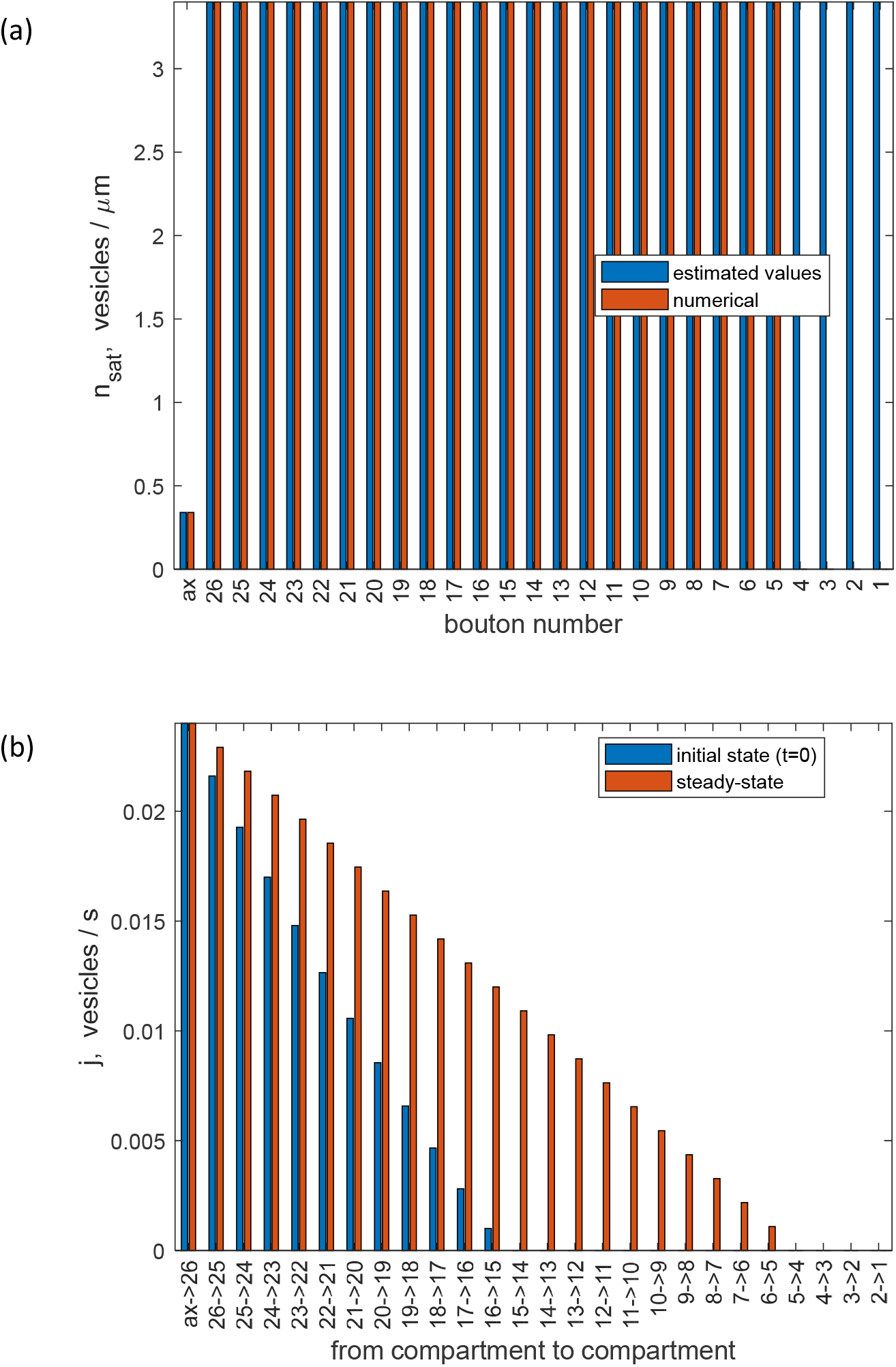
(a) Steady-state concentrations of DCVs in the resident state in various boutons and in the transiting state in the axon. Estimated values of these concentrations, calculated using Eq. (16), are compared to the values of saturated concentrations obtained numerically (at *t* →∞). Note the drop-off in the computed DCV concentrations in boutons 1 through 4. (b) Anterograde DCV fluxes from the axon to the most proximal bouton and between the boutons in the beginning of the process of filling the terminal with DCVs and at steady-state. At steady-state, anterograde fluxes between the boutons decrease until they drop to zero for the flux between boutons 5 and 4. *δ* = 0 and *n*_0,*t*_ = 1 vesicles/μm. A value of *ε* does not matter because in computing this figure it is assumed that all captured DCVs are eventually destroyed in boutons. *j*_*ax*→26_ = 2.40 ×10^−2^ vesicles/s.

It takes the DCV concentrations in the resident state in boutons 26 through 5 about 60 hours to increase to their steady-state values, with bouton 5 being the last to reach its steady-state concentration (Fig. 4). The reason why it takes longer for bouton 5 to be filled to saturation is a small anterograde flux of DCVs that comes to it from bouton 6 (Figs. 3b and 5b). Boutons 1-4 do not have any DCVs in the resident state because no anterograde DCVs reach them, they are all captured before they reach bouton 4 (Figs. 3b and 5).

**Fig. 4.**
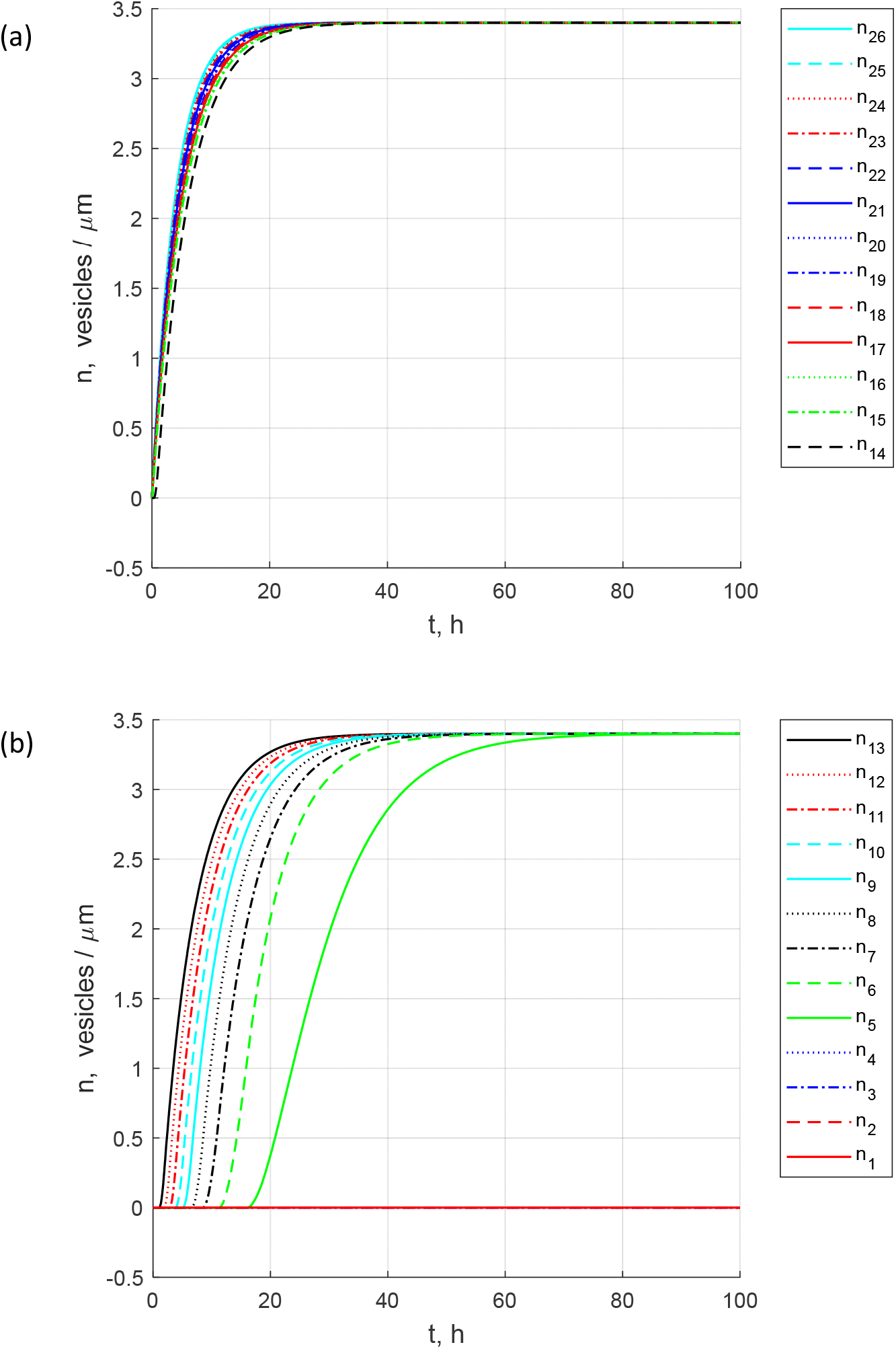
The buildup toward steady-state: concentrations of DCVs in the resident state in various boutons. (a) Boutons 26 through 14. (b) Boutons 13 through 1. Note that DCV concentrations in the four most distal boutons are equal to zero. *δ* = 0 and *n*_0,*t*_ = 1 vesicles/μm. A value of *ε* does not matter because in computing this figure it is assumed that all captured DCVs are eventually destroyed in boutons. *j*_*ax*→26_= 2.40 ×10^−2^ vesicles/s.

**Fig. 5.**
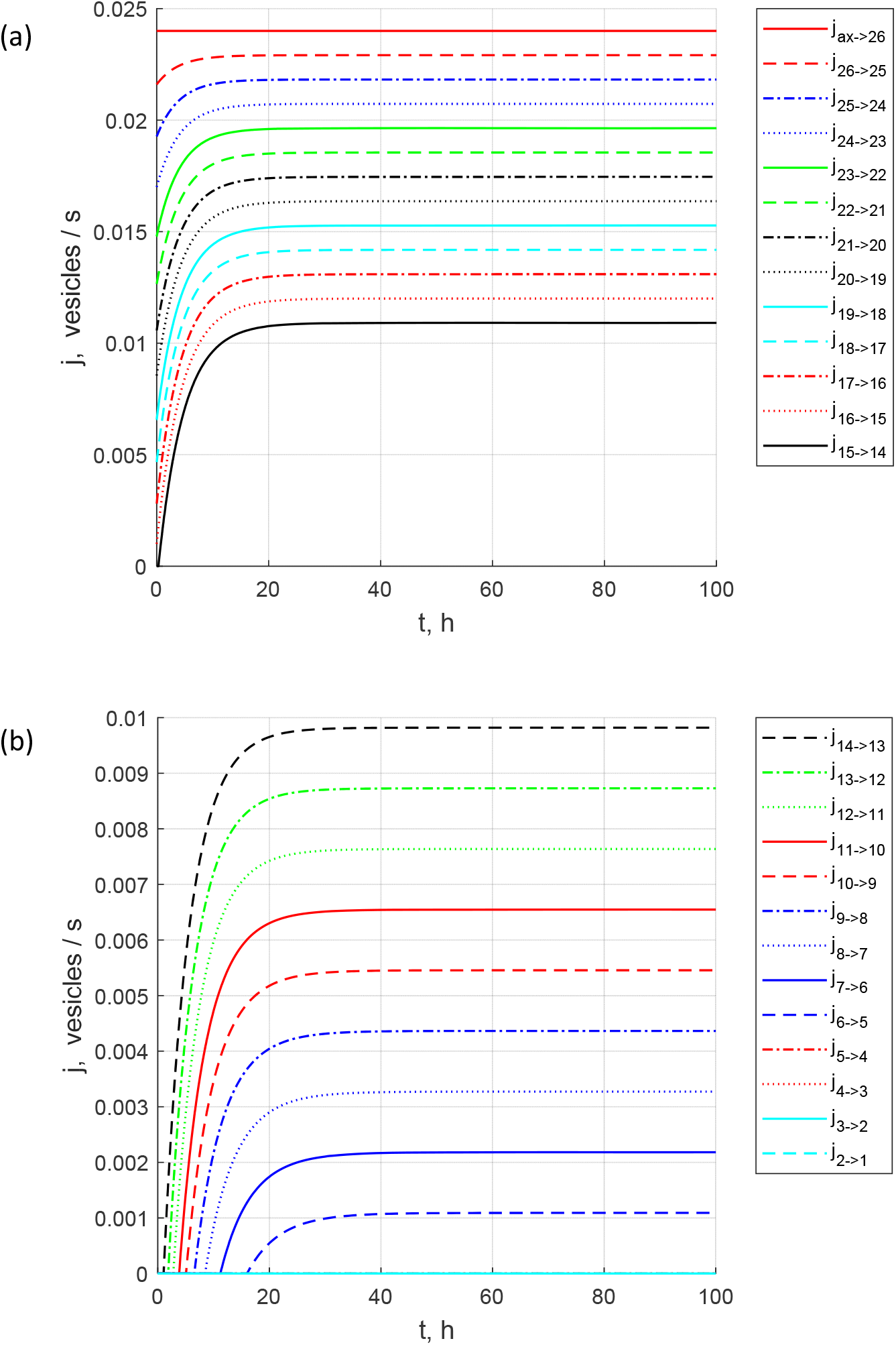
The buildup toward steady-state: the flux from the axon to the most proximal bouton and anterograde fluxes between various boutons. (a) Fluxes ax→26 through 1→14. (b) Fluxes 14→13 through 2→1. *δ* = 0 and *n*_0,*t*_ = 1 vesicles/μm. A value of *ε* does not matter because in computing this figure it is assumed that all captured DCVs are eventually destroyed in boutons. Note that the anterograde fluxes vanish after bouton 5. *j*_*ax*→26_ = 2.40 ×10^−2^ vesicles/s.

To investigate the effect of increased DCV flux from the axon into the terminal (which can be attributed to more DCV synthesis in the soma), in computing Fig. 6 we replaced Eq. (22) with the following equation:

**Fig. 6.**
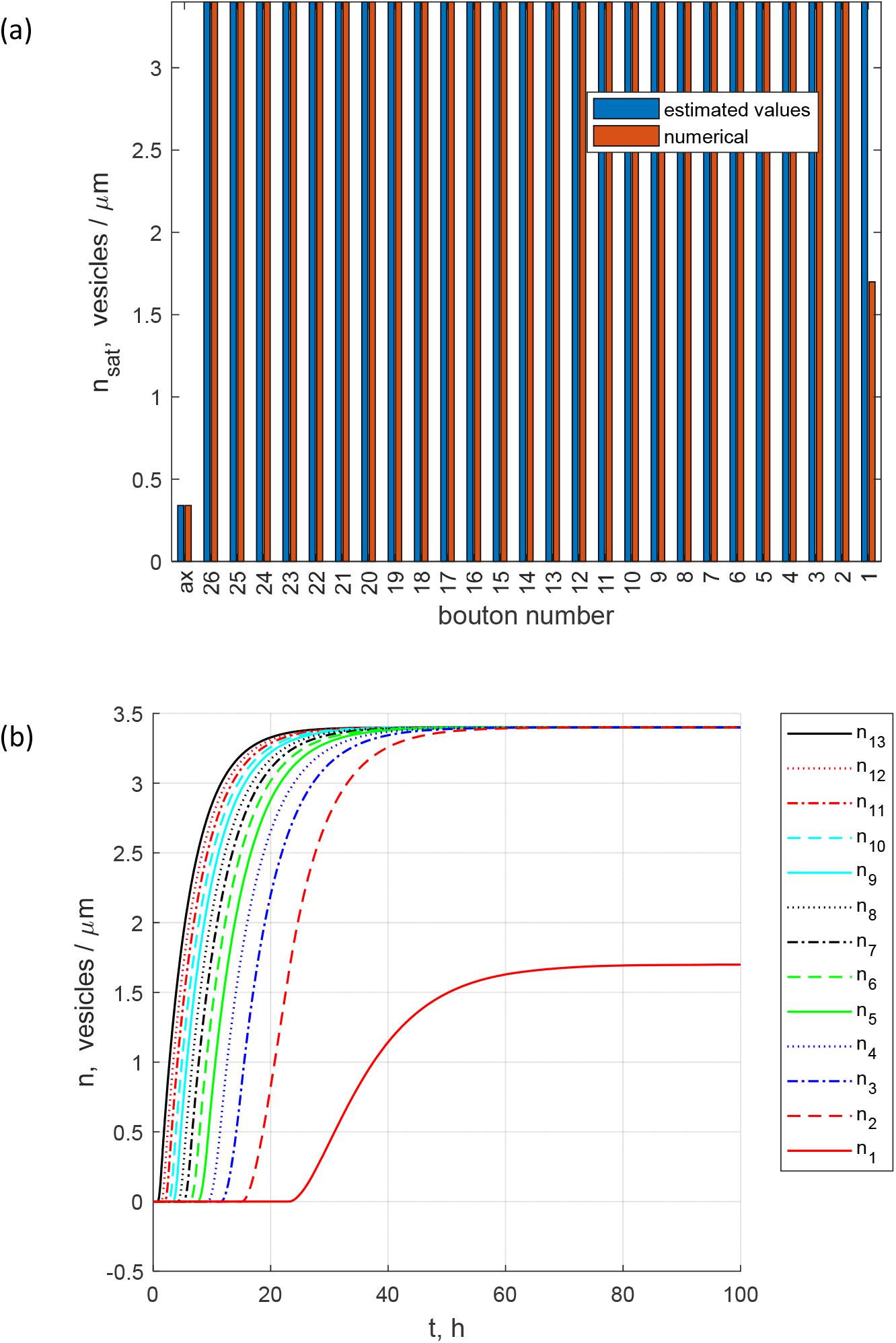
(a) Steady-state concentrations of DCVs in the resident state in various boutons and in the transiting state in the axon. Estimated values of these concentrations, calculated using Eq. (16), are compared to the values of saturated concentrations obtained numerically (at *t* →∞). Note the drop-off of the computed DCV concentration in bouton 1. (b) The buildup toward steady-state: concentrations of DCVs in the resident state in various boutons. Boutons 13 through 1. *δ* = 0 and *n*_0,*t*_ = 1 vesicles/μm. A value of *ε* does not matter because in computing this figure it is assumed that all captured DCVs are eventually destroyed in boutons. *j*_*ax*→26_= 2.78 ×10^−2^ vesicles/s.

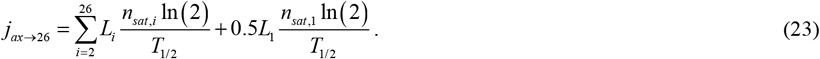

This gave *j*_*ax*→26_ = 2.78 ×10^−2^ vesicles/s, which is 15.8% greater than the value of *j*_*ax*→26_ used in computing Figs. 3-5. Bouton 1 is half-filled at steady-state (Fig. 6).

To investigate the effect of reduced DCV flux from the soma into the axon (which can be due to less DCV synthesis in the soma), in computing Fig. 7 Eq. (22) was replaced with the following equation:

**Fig. 7.**
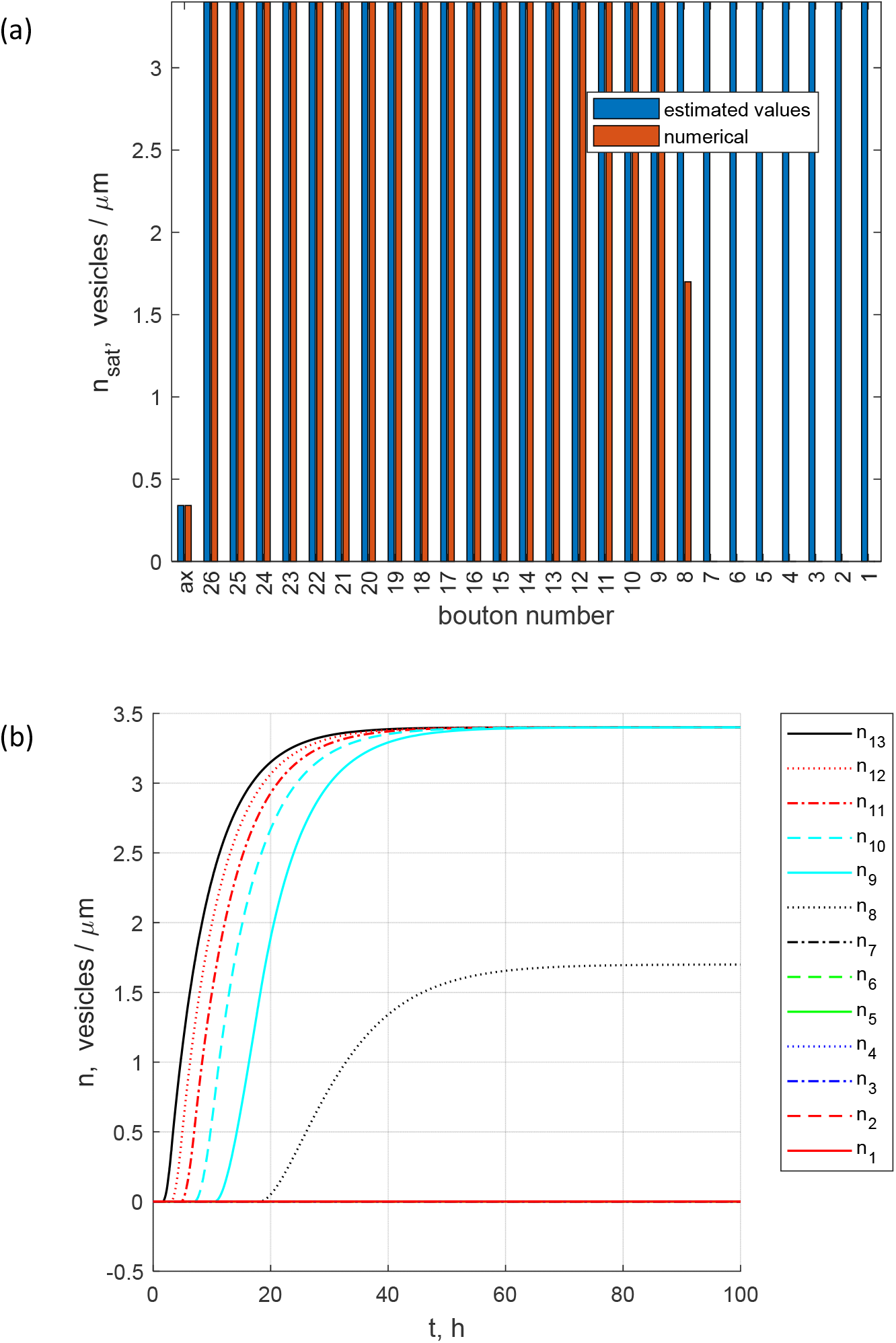
(a) Steady-state concentrations of DCVs in the resident state in various boutons and in the transiting state in the axon. Estimated values of these concentrations, calculated using Eq. (16), are compared to the values of saturated concentrations obtained numerically (at *t* →∞). Note the drop-off of the computed DCV concentration in boutons 8 through 1. (b) The buildup toward steady-state: concentrations of DCVs in the resident state in various boutons. Boutons 13 through 1. *δ* = 0 and *n*_0,*t*_ = 1 vesicles/μm. A value of *ε* does not matter because in computing this figure it is assumed that all captured DCVs are eventually destroyed in boutons. *j*_*ax*→26_ = 2.02 ×10^−2^ vesicles/s.

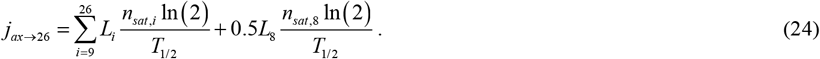

This gave *j*_*ax*→26_ = 2.02 ×10^−2^ vesicles/s, which is 15.8% smaller than the value of *j*_*ax*→26_ used in computing Figs. 3-5. At steady-state, bouton 8 is half-filled and boutons 7 through 1 are empty (Fig. 7).

Next, we investigated the situation characterized by *δ* = 0.2, *ε* = 1, and *j*_*ax*→26_ = 2.40 ×10^−2^ vesicles/s (*j*_*ax*→26_ is calculated using Eq. (22)). In this case, 20% of captured DCVs return to circulation (all of which join the anterograde component), and 80% are destroyed in the resident state in boutons. Unlike the situation displayed in Figs. 3-5, with the same value of DCV flux entering the terminal, destruction of resident DCVs in boutons at steady-state is now less due to release of some captured DCVs from the resident state. Hence, fewer DCVs need to be captured into the resident state to compensate for the DCV destruction. Also, the anterograde DCV flux is supplemented by re-released DCVs. This allows anterogradely transported DCVs to move deeper into the terminal, reaching more distal boutons (Fig. 8). The anterograde DCV flux quickly decreases from bouton to bouton. This is due to DCV capture, which is necessary to make up for the DCV loss in the resident state (Fig. 9a). However, some of the anterogradely moving DCVs reach the most distal bouton and, if they are not captured, turn around by changing the motors that propel them. The only non-zero retrograde DCV fluxes are those from bouton 1 to bouton 2 and from bouton 2 to bouton 3 (Fig. 9b).

**Fig. 8.**
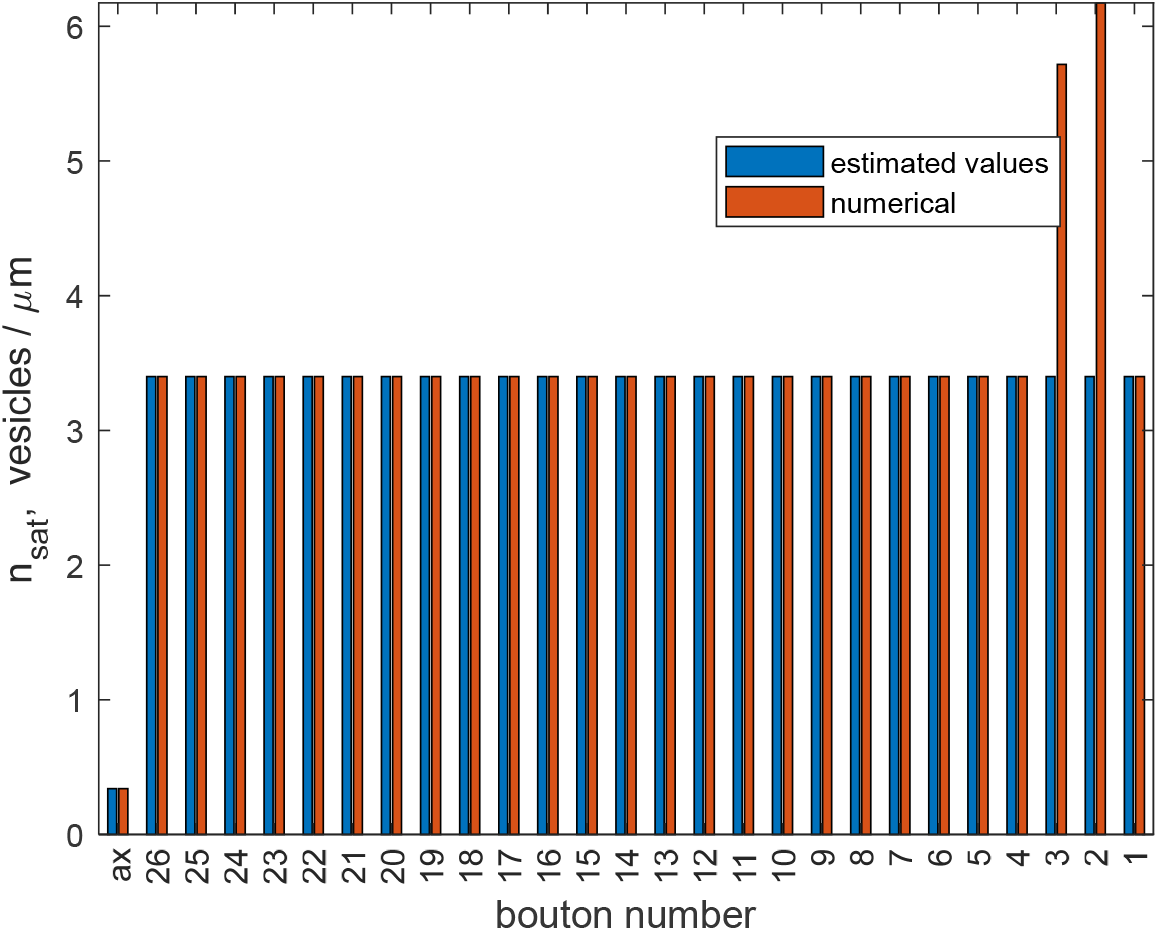
Steady-state concentrations of DCVs in the resident state in various boutons and in the transiting state in the axon. Estimated values of these concentrations, calculated using Eq. (16), are compared to the values of saturated concentrations obtained numerically (at *t* →∞). Note the increased concentration of DCVs predicted by the model in boutons 1 and 2. *δ* = 0.1, *n*_0,*t*_ = 1 vesicles/μm, and *ε* = 1. *j*_*ax*→26_= 2.40 ×10^−2^ vesicles/s.

**Fig. 9.**
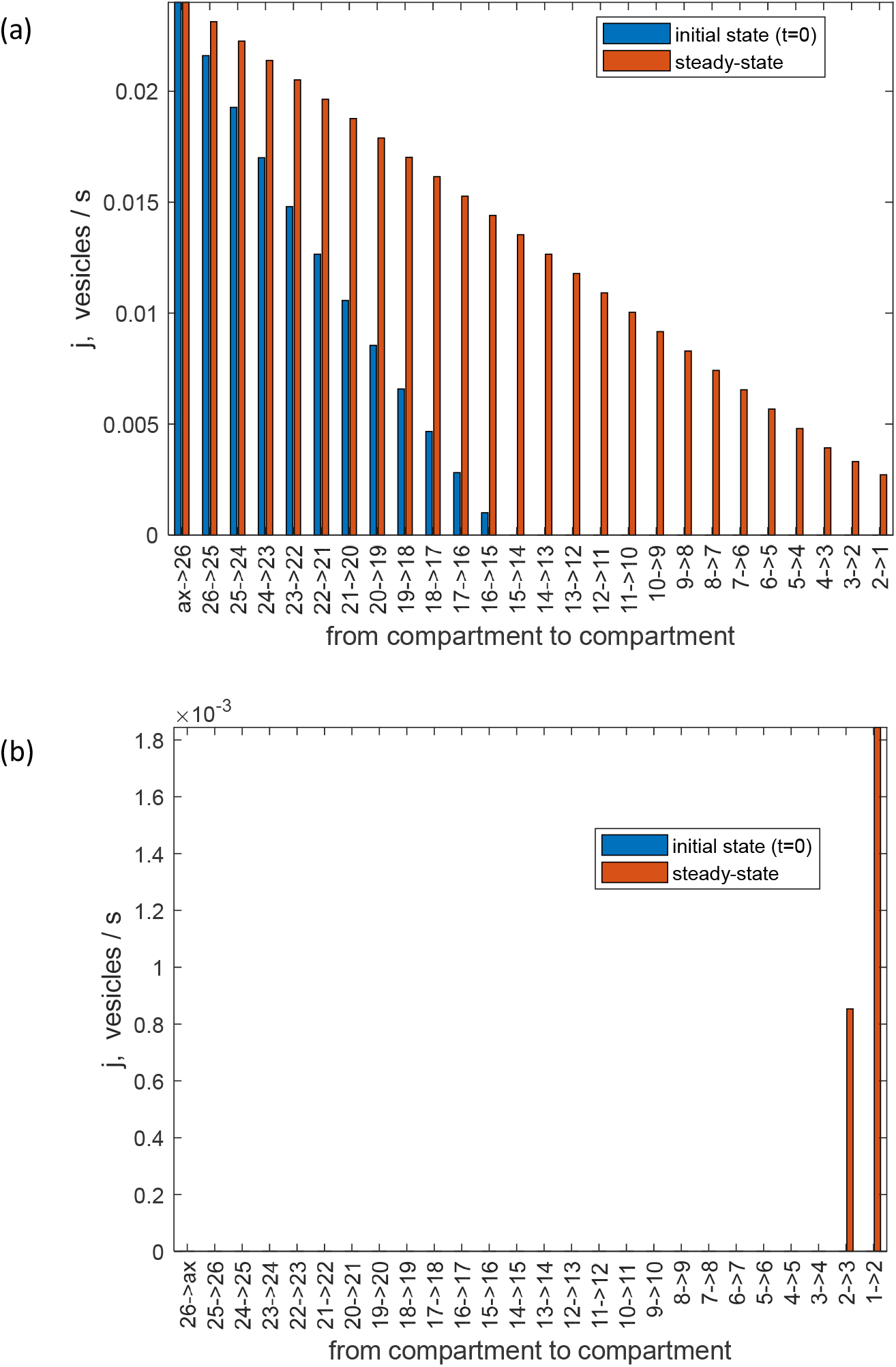
(a) Anterograde DCV fluxes from the axon to the most proximal bouton and between the boutons at the beginning of the process of filling the terminal with DCVs and at steady-state. At steady-state, anterograde fluxes between the boutons decrease but reach bouton 1. (b) Retrograde DCV fluxes between the boutons in the beginning of the process of filling the terminal with DCVs and at steady-state. The only non-zero retrograde flux is that from bouton 1 to bouton 2. *δ* = 0.2, *n*_0,*t*_ = 1 vesicles/μm, and *ε* = 1. *j*_*ax*→26_ = 2.40 ×10^−2^ vesicles/s.

It takes more time for the DCV concentrations in more distal boutons to reach their steady-state values. This applies to boutons 13 through 3, but especially to boutons 3 and 2, which capture not only anterograde but also retrograde DCVs. It takes approximately 77.8 hours for the DCV concentration in bouton 3 to reach its steady-state value (Fig. 10b, Table 4). We note that the duration of *Drosophila* third instar (used in the experiments reported in [18]) is approximately 48 hours, meaning that steady-state may not be reached in *Drosophila* terminals.

**Table 4.**
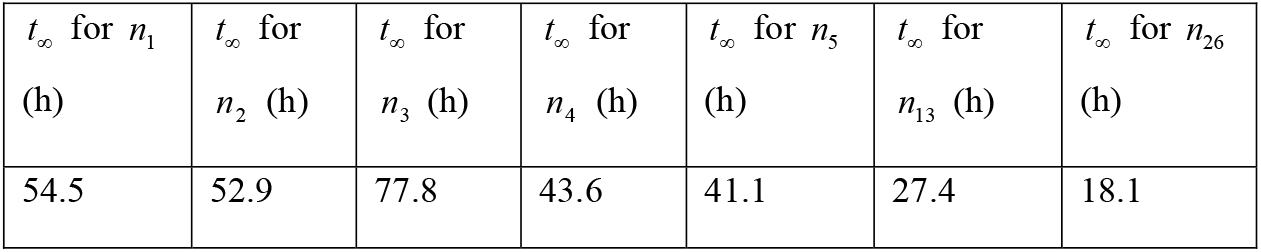
Approximate time, *t*_∞_, needed for resident DCVs in seven representative boutons (#1, 2, 3, 4, 5, 13, and 26) to reach steady-state concentrations. We assumed that when the DCV concentration in a bouton exceeds 99% of the steady-state concentration in the corresponding bouton, the steady-state is reached. *δ* = 0.2, *n*_0,*t*_ =1, and *ε* = 1. *j*_*ax*→26_ = 2.40 ×10^−2^ vesicles/s.

**Fig. 10.**
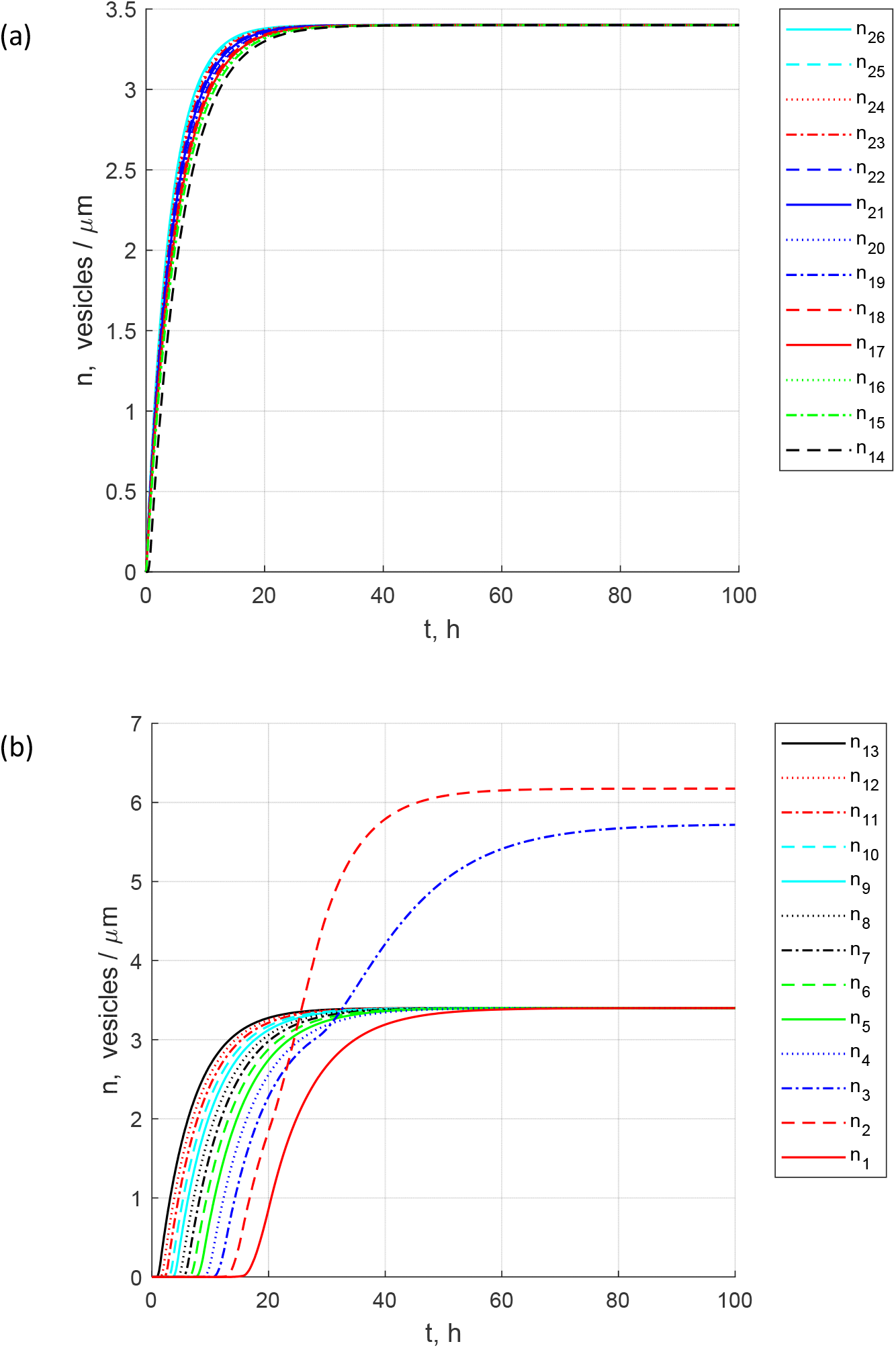
The buildup toward steady-state: concentrations of DCVs in the resident state in various boutons. (a) Boutons 26 through 14. (b) Boutons 13 through 1. Note the increased DCV concentrations in boutons 2 and 1. *δ* = 0.2, *n*_0,*t*_ = 1 vesicles/μm, and *ε* = 1. *j*_*ax*→26_ = 2.40 ×10^−2^ vesicles/s.

The above observations point to an interesting phenomenon. Since retrogradely moving DCVs are also captured in boutons, the concentration of resident DCVs in bouton 2 is 82% larger than in other boutons and in bouton 3 is 68% larger than in other boutons. Thus, the model predicts that if DCVs are re-released from boutons, the concentration in most distal boutons (except bouton 1) is likely elevated. Note that the DCV concentration in bouton 1 is not elevated because, according to our model, DCVs pass bouton 1 only once, and thus there is no “double capture” of anterogradely and then retrogradely moving DCVs in bouton 1.

Anterograde fluxes between the boutons decay quickly toward more distal boutons because of DCV capture and destruction in more proximal boutons (Fig. 11). The only non-zero retrograde DCV fluxes between the boutons are those from bouton 1 to bouton 2 and from bouton 2 to bouton 3 (Fig. 12a).

**Fig. 11.**
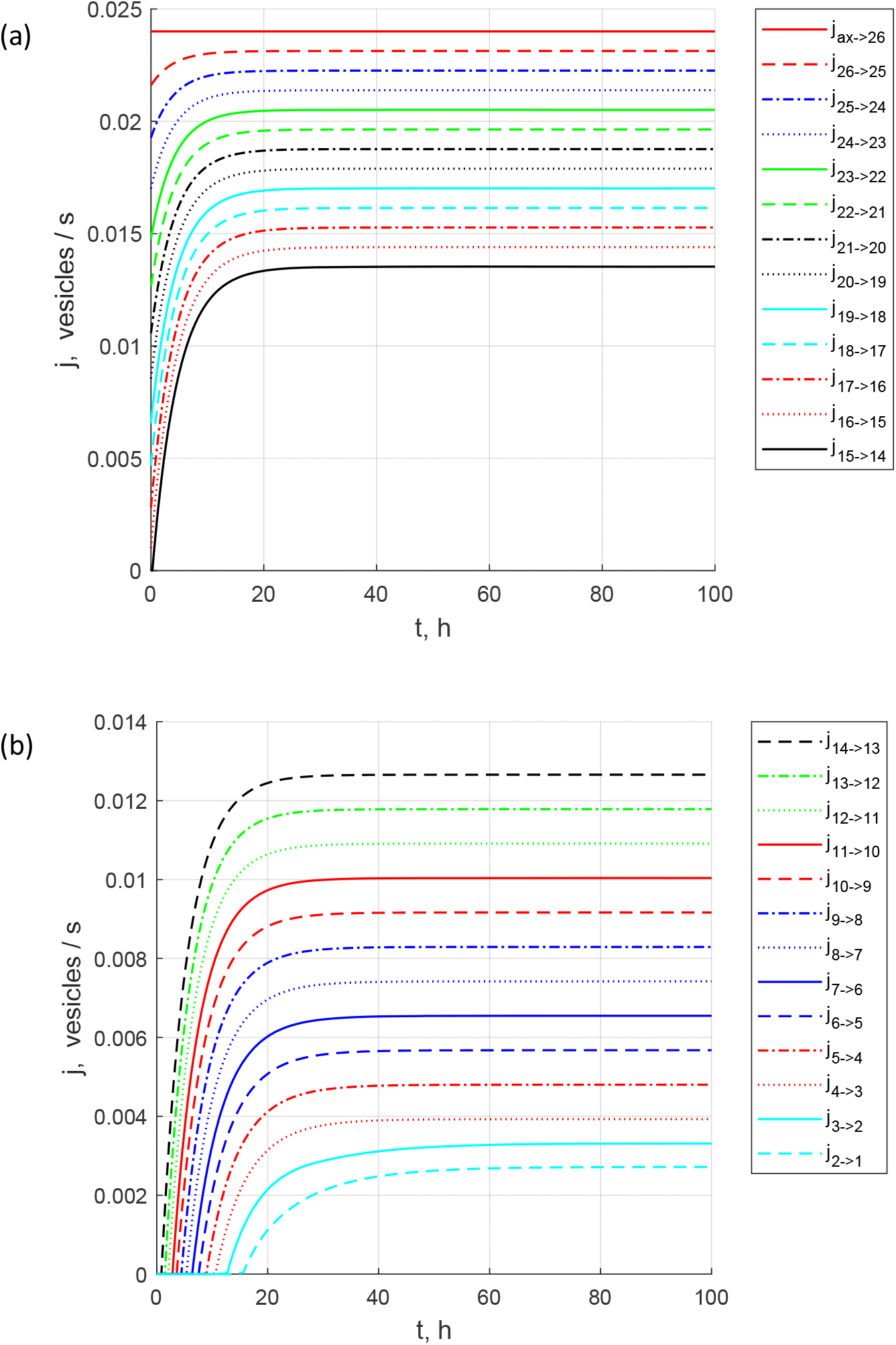
The buildup toward steady-state: the flux from the axon to the most proximal bouton and anterograde fluxes between various boutons. (a) Fluxes ax→26 through 1→14. (b) Fluxes 14→13 through 2→1. *δ* = 0.2, *n*_0,*t*_ = 1 vesicles/μm, and *ε* = 1. *j*_*ax*→26_ = 2.40 ×10^−2^ vesicles/s.

**Fig. 12.**
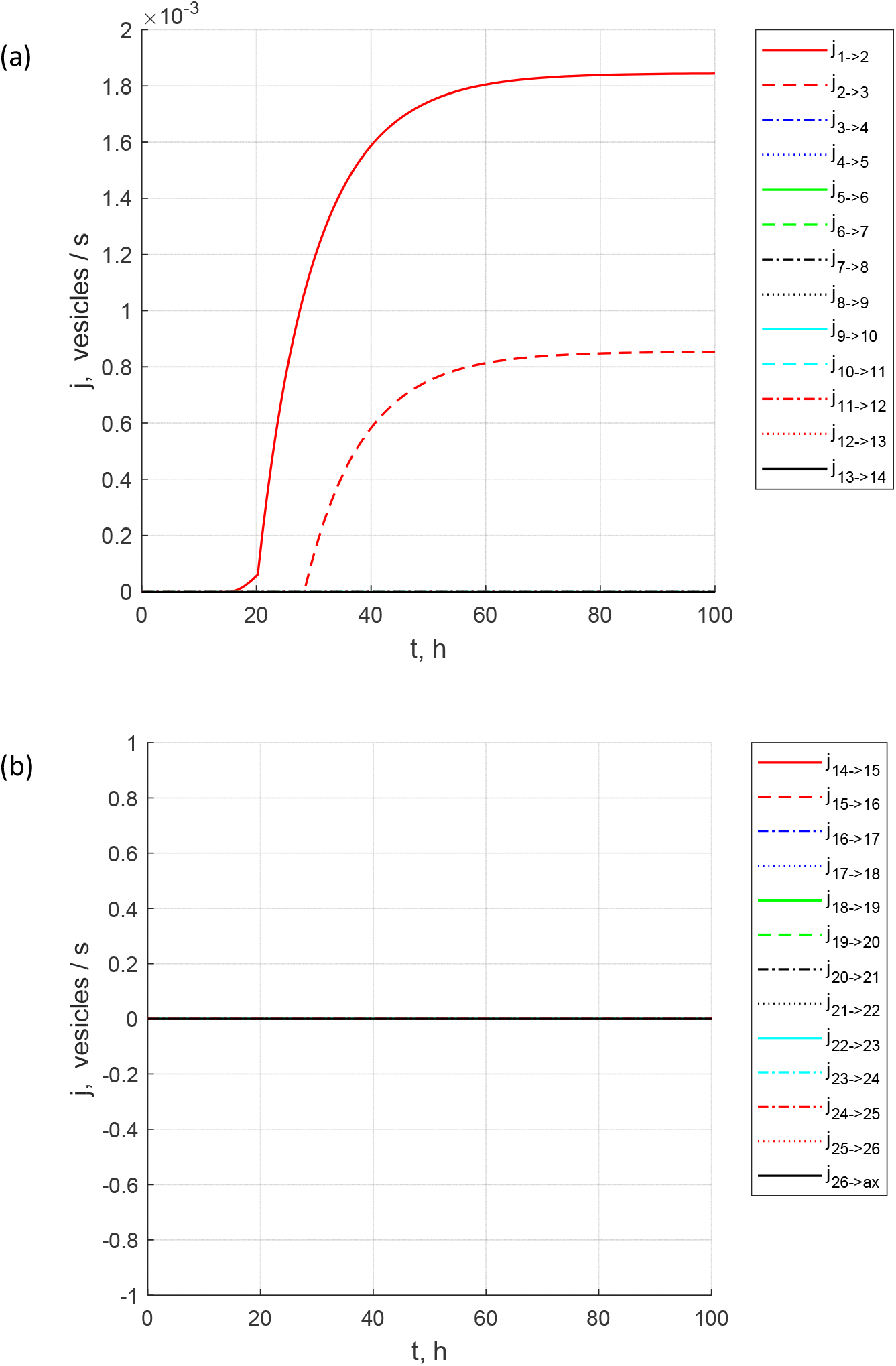
The buildup toward steady-state: the flux from the axon to the most proximal bouton and anterograde fluxes between various boutons. (a) Fluxes 1→2 through 13→14. (b) Fluxes 14→15 through 26→ax. Note that the only non-zero retrograde fluxes are from bouton 1 to bouton 2 and from bouton 2 to bouton 3. *δ* = 0.2, *n*_0,*t*_ = 1 vesicles/μm, and *ε* = 1. *j*_*ax*→26_ = 2.40 ×10^−2^ vesicles/s.

To investigate the effect of increased DCV flux from the soma into the axon (which may be due to more DCV synthesis in the soma), in computing Figs. 13 and 14 we replaced Eq. (22) with the following equation:

**Fig. 13.**
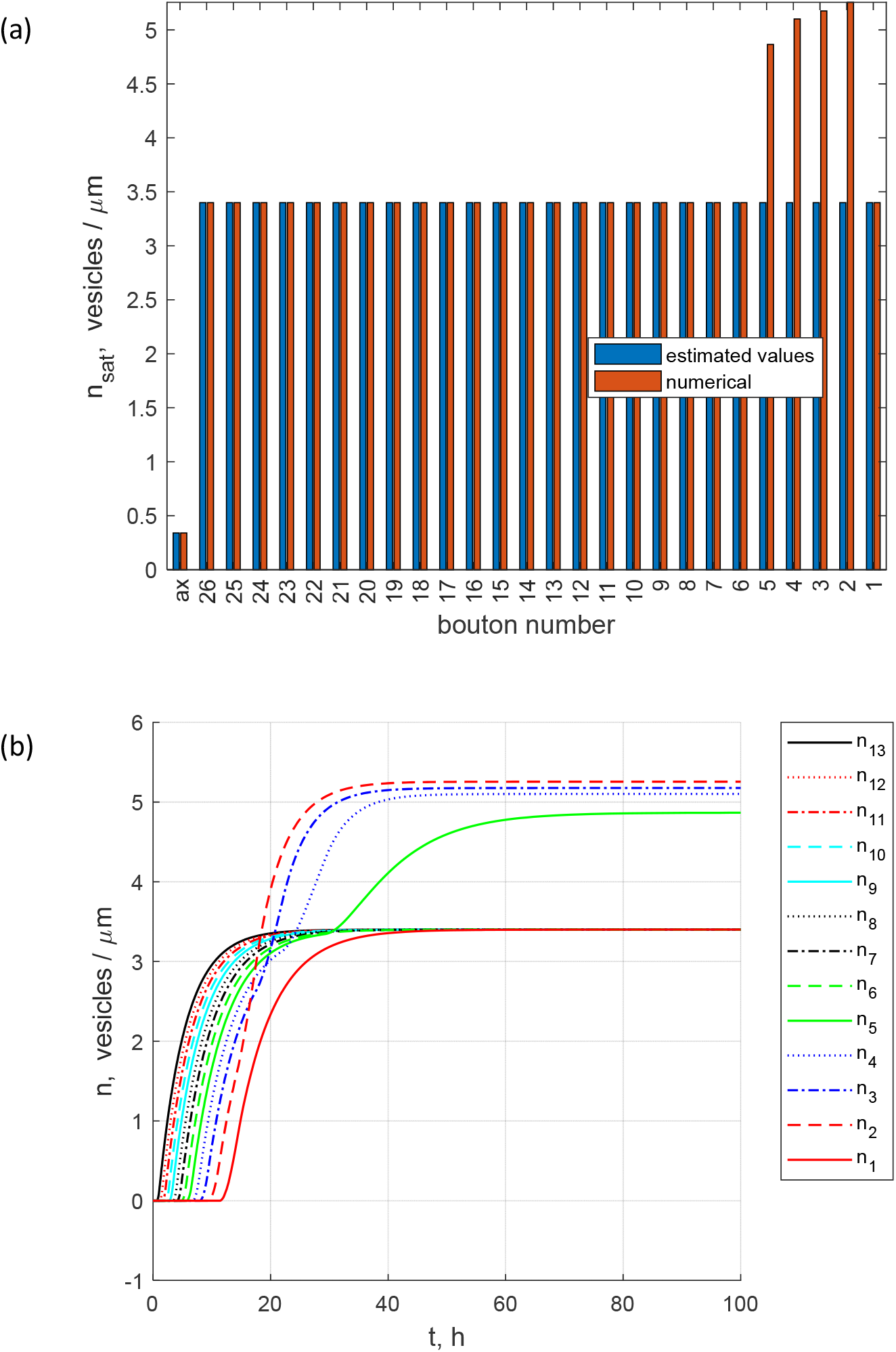
(a) Steady-state concentrations of DCVs in the resident state in various boutons and in the transiting state in the axon. Estimated values of these concentrations, calculated using Eq. (16), are compared to the values of saturated concentrations obtained numerically (at *t* →∞). (b) The buildup toward steady-state: concentrations of DCVs in the resident state in various boutons. Boutons 13 through 1. Note the increased DCV concentrations in boutons 2, 3, 4 and 5. *δ* = 0 and *n*_0,*t*_ = 1 vesicles/μm. A value of *ε* does not matter because all captured DCVs are eventually destroyed in boutons. *j*_*ax*→26_= 3.05×10^−2^ vesicles/s.

**Fig. 14.**
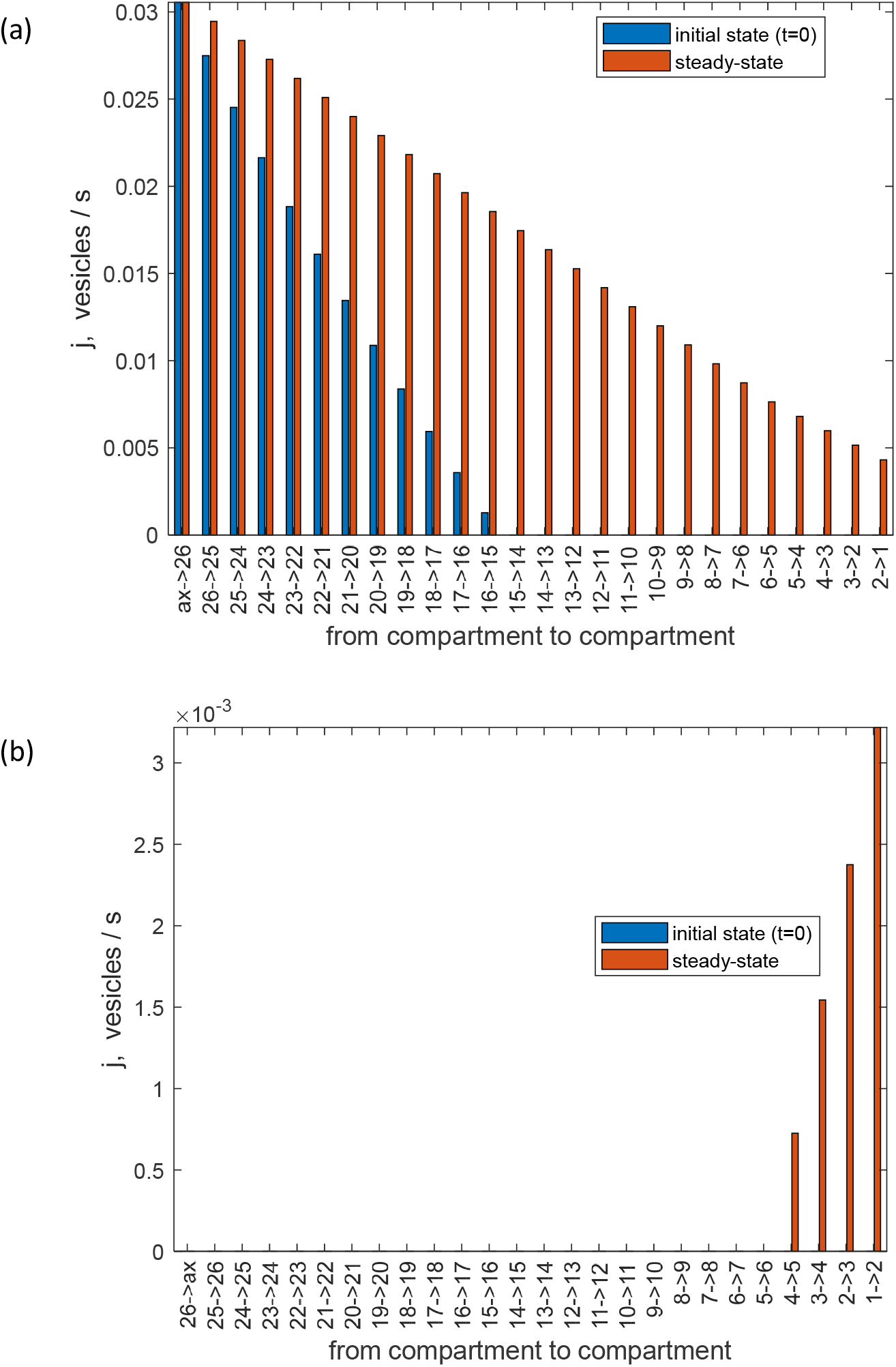
(a) Anterograde DCV fluxes from the axon to the most proximal bouton and between the boutons at the beginning of the process of filling the terminal with DCVs and at steady-state. At steady-state, anterograde fluxes between the boutons decrease but reach bouton 1. (b) Retrograde DCV fluxes between the boutons in the beginning of the process of filling the terminal with DCVs and at steady-state. The only non-zero retrograde fluxes are those from bouton 1 to bouton 2, 2 to 3, 3 to 4, and 4 to 5. *δ* = 0 and *n*_0,*t*_ = 1 vesicles/μm. A value of *ε* does not matter because all captured DCVs are eventually destroyed in boutons. *j*_*ax*→26_= 3.05×10^−2^ vesicles/s.

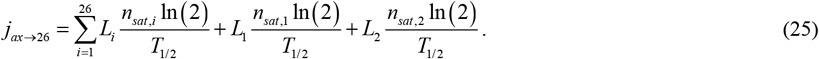

This gave *j*_*ax*→26_ = 3.05×10^−2^ vesicles/s, which is 27.1% greater than the value of *j*_*ax*→26_ used in computing Figs. 3-5. Boutons 2 through 5 have increased DCV content at steady-state (Fig. 13). Anterograde fluxes slowly decrease from the proximal to distal boutons, and as DCVs switch to retrograde motion in the most distal bouton, the fluxes continue to decrease until they drop to zero between boutons 5 and 6 (Fig. 14).

## 4. Discussion, limitations of the model and future directions

The contribution of this paper is that it establishes that the zone with a sudden drop-off of the DCV content at the end of the terminal, which was reported in [18], can be explained without assuming any special properties for the distal boutons. The drop-off is caused by the depletion of the anterograde flux of DCVs due to DCV capture in more proximal boutons as DCVs pass them rather than by the inability of distal boutons to capture more DCVs. The obtained results suggest that the size of the sudden drop-off zone in the DCV content depends on the anterograde DCV flux entering the terminal.

Our modeling results suggest that to explain the sudden drop-off in the DCV content in distal boutons, the assumption that captured DCVs and their content are destroyed in boutons rather than re-released to the circulation is needed. Furthermore, the model predicts that an increase of the DCV anterograde flux in the terminal can lead to the disappearance of the drop-off zone. A further increase of the flux leads to DCVs turning around at the most distal bouton and moving retrogradely. As this happens, DCVs can be captured again in boutons which they have passed. If all of the retrogradely moving DCVs are captured in distal boutons, this could result in an increase of the DCV concentration in the distal boutons. This prediction is both testable and falsifiable. The model suggests that a 27% increase in the anterograde DCV flux coming from the soma can make a difference between a drop-off zone and a zone with an increased DCV content in distal boutons.

The main limitation of the model is the simplification of the process of DCV capture and release by the boutons. Future research should address the situation when DCVs release some of their contents by the kiss and run exocytosis, in contrast to traditional full collapse exocytosis, which fully empties DCVs [33]. Theoretically, the compartmental model can be replaced with a more elegant partial differential equation (PDE), which would collapse to the present model upon discretization. This approach is attractive because the obtained PDE could be investigated for instability, leading to predicting a precise location of the drop-off zone. An alternative approach could be based on developing a discrete stochastic simulation model, which would treat DCVs as individual vesicles.

## Acknowledgment

IAK acknowledges the fellowship support of the Paul and Daisy Soros Fellowship for New Americans and the NIH/National Institutes of Mental Health (NIMH) Ruth L. Kirchstein NRSA (F30 MH122076-01). AVK acknowledges the support of the National Science Foundation (award CBET-2042834) and the Alexander von Humboldt Foundation through the Humboldt Research Award.

